# TNPO3-mediated nuclear entry of the Rous sarcoma virus Gag protein is independent of the cargo-binding domain

**DOI:** 10.1101/2020.03.12.989608

**Authors:** Breanna L. Rice, Matthew S. Stake, Leslie J. Parent

**Affiliations:** Division of Infectious Diseases and Epidemiology, Department of Medicine, Penn State College of Medicine, Hershey, PA, USA; Department of Microbiology and Immunology, Penn State College of Medicine, Hershey, PA, USA

**Author notes:** Address correspondence to Leslie Parent,. B.L.R and M.S.S. contributed equally to this work.

## Abstract

Retroviral Gag polyproteins orchestrate the assembly and release of nascent virus particles from the plasma membranes of infected cells. Although it was traditionally thought that Gag proteins trafficked directly from the cytosol to the plasma membrane, we discovered that the oncogenic avian alpharetrovirus Rous sarcoma virus (RSV) Gag protein undergoes transient nucleocytoplasmic transport as an intrinsic step in virus assembly. Using a genetic approach in yeast, we identified three karyopherins that engage the two independent nuclear localization signals (NLS) in Gag. The primary NLS is in the nucleocapsid (NC) domain of Gag and binds directly to importin-α, which recruits importin-β to mediate nuclear entry. The second NLS, which resides in the matrix (MA) domain, is dependent on importin-11 and transportin-3 (TNPO3), known as MTR10p and Kap120p in yeast, although it is not clear whether these import factors are independent or additive. The functionality of importin α/β and importin-11 has been verified in avian cells, whereas the role of TNPO3 has not been studied. In this report, we demonstrate that TNPO3 mediates nuclear entry of Gag and directly binds to Gag. To our surprise, this interaction did not require the cargo-binding domain of TNPO3, which typically mediates nuclear entry for other binding partners of TNPO3 including SR-domain containing splicing factors and tRNAs that re-enter the nucleus. These results suggest that RSV hijacks the host nuclear import pathway using a unique mechanism, potentially allowing other cargo to bind TNPO3 simultaneously.

**Importance:** RSV Gag nuclear entry is facilitated using three distinct host import factors that interact with nuclear localization signals in the Gag MA and NC domains. Here we show that the MA region is required for nuclear import of Gag through the TNPO3 pathway. Gag nuclear entry does not require the cargo binding domain of TNPO3. Understanding the molecular basis for TNPO3-mediated nuclear trafficking of the RSV Gag protein may lead to a deeper appreciation for whether different import factors play distinct roles in retrovirus replication.

## Introduction

The retrovirus structural protein Gag is a multi-domain protein that is responsible for packaging the viral genome and directing the assembly and budding of virus particles from the plasma membrane of infected cells. The MA (matrix) domain of Gag facilitates membrane targeting and binding. The CA (capsid) domain is important for Gag protein-protein interactions, as well forming the virus particle capsid. The NC (nucleocapsid) binds to the viral RNA genome (vRNA) for packaging and is involved in protein-protein interactions that promote muiltimerization and virus assembly (34, 53).

It has been found that the Gag proteins from various retroviruses undergo nuclear trafficking [reviewed in (49)]. The mechanisms of Gag nuclear trafficking are not completely understood. For Rous sarcoma virus (RSV), it is hypothesized that the reason for Gag nuclear trafficking is binding vRNA for packaging (13). The nuclear export of RSV Gag was first discovered to be dependent on the CRM1 nuclear export protein, which interacts with a nuclear export signal (NES) mapped to the p10 domain (Figure 1A) (42, 44). The MA and NC domains were later found to be involved in the nuclear import of Gag through their NLSs. Studies utilizing *Saccharomyces cerevisiae* mutants deficient in members of the Importin-β protein superfamily found that the NC domain undergoes nuclear entry through the Kap60p/Kap95p (importin-α/β) pathway, while MA uses either Kap120p (importin 11) or Mtr10p, also known as transportin-SR, transportin 3 or TNPO3 in higher eukaryotes (5). The interactions between the NC domain and the importin-α/β (Impα/β) complex, as well as the MA domain and importin 11 (Imp11) were confirmed via affinity-tagged purifications (14).

**Figure 1:**
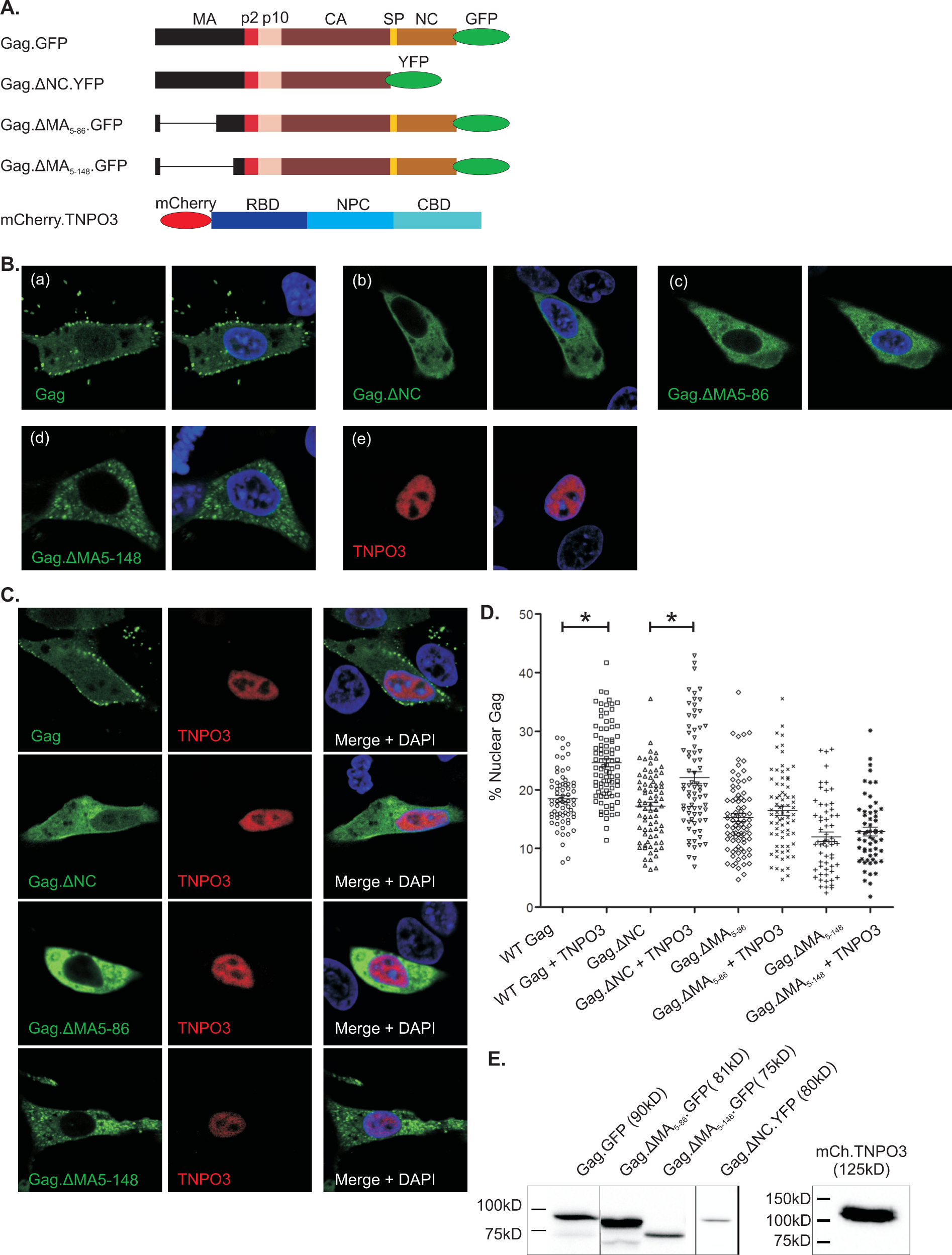
Effects of TNPO3 expression on nuclear accumulation of Gag. (A) Schematic representation of RSV Gag constructs used, including the Gag domains MA (matrix), p2, p10, CA (capsid), SP (spacer), and NC (nucleocapsid) with an in frame fusion of GFP. Gag deletion mutants have YFPor GFP fusions, as indicated. TNPO3 consists of a three domain structure including an N-terminal RanGTP binding domain (RBD), a nuclear pore complex (NPC) binding domain, and C-terminal cargo-binding domain (CBD). (B) The localization of wildtype Gag.GFP (a) Gag.ΔNC.YFP (b), Gag.ΔMA_5-86_.GFP (c) and Gag.ΔMA_5-148_.GFP (d) expressed in QT6 cells are shown with DAPI-stained images included to show the nucleus of the cell. Panel (e) shows the localization of mCherry.TNPO3. (C) Cells co-expressing the Gag wild-type or mutant proteins with mCherry.TNPO3 are shown. (D) A scatter plot showing the percentage of Gag localized to the nucleus (% nuclear compared to the signal in the entire cell) for each cell analyzed, with or without co-expression of mCherry.TNPO3. At least 60 cells were analyzed from three independent experiments for each condition, with the standard error of the mean represented by the error bars. Statistical analysis was performed using unpaired Student’s t-test and significance (p < 0.0001) is indicted by an asterisk. In cases where mCherry.TNPO3 and Gag constructs were co-transfected, only cells expressing both Gag and mCherry.TNPO3 were analyzed quantitatively and shown in the graph. Images were chosen for display that are representative of the mean fluorescence intensity of the population analyzed. (E) Western blot of cell lysates transfected with the indicated plasmids were performed using anti-GFP or α-mCherry antibodies, as appropriate, to show that full-length proteins were expressed at the expected sizes with minimal proteolysis for Gag.GFP, Gag variants, and mCherry.TNPO3. The vertical line between Gag.GFP and Gag. ΔMA_5-86_ .GFP represents the removal of an irrelavant lane in the gel. Gag.ΔNC.YFP was expressed on a separate blot, as was mCherry.TNPO3 (mCh.TPO3).

TNPO3 is a member of the Importin-β family of karyopherins (24, 29) and serves as a nuclear import receptor for a class of evolutionarily conserved, essential splicing factors and pre-mRNA processing proteins known as serine and arginine-rich (SR) proteins. SR proteins derive their names from the serine/arginine enriched motifs in their C-terminal regions, and they contain RNA recognition domains located at their N termini (47, 54). The RS domain of splicing factors interact directly with the C-terminal CBD of TNPO3, mediating nuclear entry of the splicing factor (17, 23, 31). TNPO3 also mediates nuclear import of the pre-mRNA splicing factor RBM4 through its interaction with stretches of alanine residues, termed polyalanine domains (22) and has been implicated in mediating nucleocytoplasmic tRNA transport in yeast and human cells (35, 46, 55).

TNPO3 has also been shown to be important for the replication of several retroviruses, including human immunodeficiency virus type 1 (HIV-1) (4, 7, 19). TNPO3 is involved in early events of HIV infection, primarily at the level of pre-integration complex (PIC) nuclear entry (55). *In vitro* and *in vivo* studies have shown direct interactions between TNPO3 and HIV integrase (9, 25, 27, 30, 51). In cells depleted of TNPO3 by small hairpin RNA (shRNA), HIV-1 PIC nuclear entry is impaired and integration is reduced (11, 45, 52). Further, the cargo-binding domain of TNPO3 is required to rescue HIV infection in cells depleted of TNPO3 by shRNA (29). Proviruses that do manage to integrate in TNPO3 knockdown cells do so in regions with lower gene density than the integration sites in control cells (38, 41). There is also evidence that HIV CA governs TNPO3 sensitivity. Amino acid substitutions in CA, such as the N74D mutation, permit PIC nuclear entry in TNPO3 knockdown cells through an altered import pathway (10, 26, 45, 55). The actual mechanism by which TNPO3 depletion impairs HIV infectivity is not well understood but is thought to involve the cleavage and polyadenylation specificity factor subunit 6 (CPSF6), an SR protein (3, 11, 12, 16, 39).Other studies have demonstrated that TNPO3 is important for the nuclear import of the foamy virus (FV) PIC. When TNPO3 expression is decreased in cells, the concentration of FV integrase in the nucleus was reduced, although the nuclear localization of FV Gag was not affected (2). TNPO3 has also been shown to be important during infection of simian immunodeficiency virus mac239, equine infectious anemia virus (21, 29), HIV-2 (7), and bovine immunodeficiency virus (21). Although it appears that TNPO3 is important for a number of retroviruses during early infection, the interaction of TNPO3 with the Gag polyprotein has yet to be investigated in depth and is the subject of this work.

## Results

Previous studies examining the role of host importins in RSV Gag nuclear import identified three import factors: Impβ, Imp11, and TNPO3 (5). Imp11 binds directly to the MA domain of Gag whereas the and Impβ protein binds to the NC NLS and recruits Impα to mediate Gag nuclear entry. The interaction of RSV Gag with TNPO3 was not further explored until this time (14).

The domain organization of the RSV Gag polyprotein, which consists of MA, p2, p10, CA, SP, and NC is shown in Figure 1A. TNPO3 consists of three domains: the RanGTP binding domain (RBD), the nuclear pore complex (NPC) interaction domain, and the cargo binding domain (CBD) (Fig. 1A). RSV Gag.GFP is normally distributed in the nucleus, cytoplasm, and along the plasma membrane of avian cells [Fig. 1B; (13, 18, 36, 42-44)]. Expression of mCherry.TNPO3 resulted in an increase in the amount of Gag protein located within the nucleus (Fig. 1C, D). Quantitation of the amount of nuclear Gag fluorescence demonstrated an increase from 18.5% to 25% with co-expression of mCherry.TNPO3 (p < 0.0001) (Fig. 1D). Cells representative of the mean fluorescence intensity of the population of cells analyzed are shown in Figure 1B and C.

Based on the finding that RSV MA nuclear entry depends on TNPO3/Mtr10p in *Saccharyomyces cerevisae* (5), we examined whether this interaction was functionally relevant in avian cells. To this end, two mutants were utilized that contain deletions in MA (Fig. 1A). One deletion mutant is missing the N-terminal portion of MA containing the previously-identified NLS (Gag.ΔMA_5-86_), and the other has a large internal deletion of MA extending from residues 5-148 (Gag.ΔMA_5-148_). Nuclear localization of both MA mutants, Gag.ΔMA_5-86_ and Gag.ΔMA_5-148_, was slightly reduced compared to full-length Gag (18.5%), with quantitative analysis showing that 15.5% and 12% of the mutant Gag proteins were nuclear localized, respectively (Fig. 1B, D). When mCherry.TNPO3 was co-expressed, the amount of nuclear Gag did not change significantly (16.5% and 13% for Gag.ΔMA_5-86_ and Gag.ΔMA_5-148_, respectively) (Fig. 1C, D). These results suggested that the activity of TNPO3 in mediating nuclear entry of Gag is dependent on the MA domain.

By contrast, when the NC domain of Gag was deleted (Gag.ΔNC, Fig. 1A), a result similar to wild-type Gag was observed with co-expression of mCherry.TNPO3, with an increase in the baseline nuclear localization of Gag.ΔNC, from 17% to 22% (p < 0.0001; Fig. 1B-D). This result was anticipated based on our previous results indicating that nuclear import of RSV NC is mediated by the importin α/β complex (5, 14) but not TNPO3, and validated our previous findings. To examine whether the Gag constructs expressed in cells produced full-length proteins, SDS-PAGE and western blot analysis was performed of the lysates expressing each Gag protein (Figure 1E), demonstrating that all of the constructs were of the expected molecular weight.

The next goal was to identify which domain of TNPO3 was required for Gag nuclear import in avian cells. We anticipated that the cargo-binding domain (CBD) would be necessary because of previous findings demonstrating SR proteins bind directly to the CBD for nuclear import (17, 23). To test this hypothesis, the mCherry.TNPO3.ΔCargo mutant was co-expressed in cells with Gag.YFP (Fig. 2A). Unexpectedly, there was an increase in the amount of nuclear Gag from 19% to 26% (p < 0.0001) (Fig. 2B, C), indicating the the TNPO3 CBD was dispensable for Gag nuclear entry. To examine Gag.YFP and mCherry.TNPO3.ΔCargo proteins expressed in cells, SDS-PAGE and western blot analysis were performed (Fig 2D).

**Figure 2:**
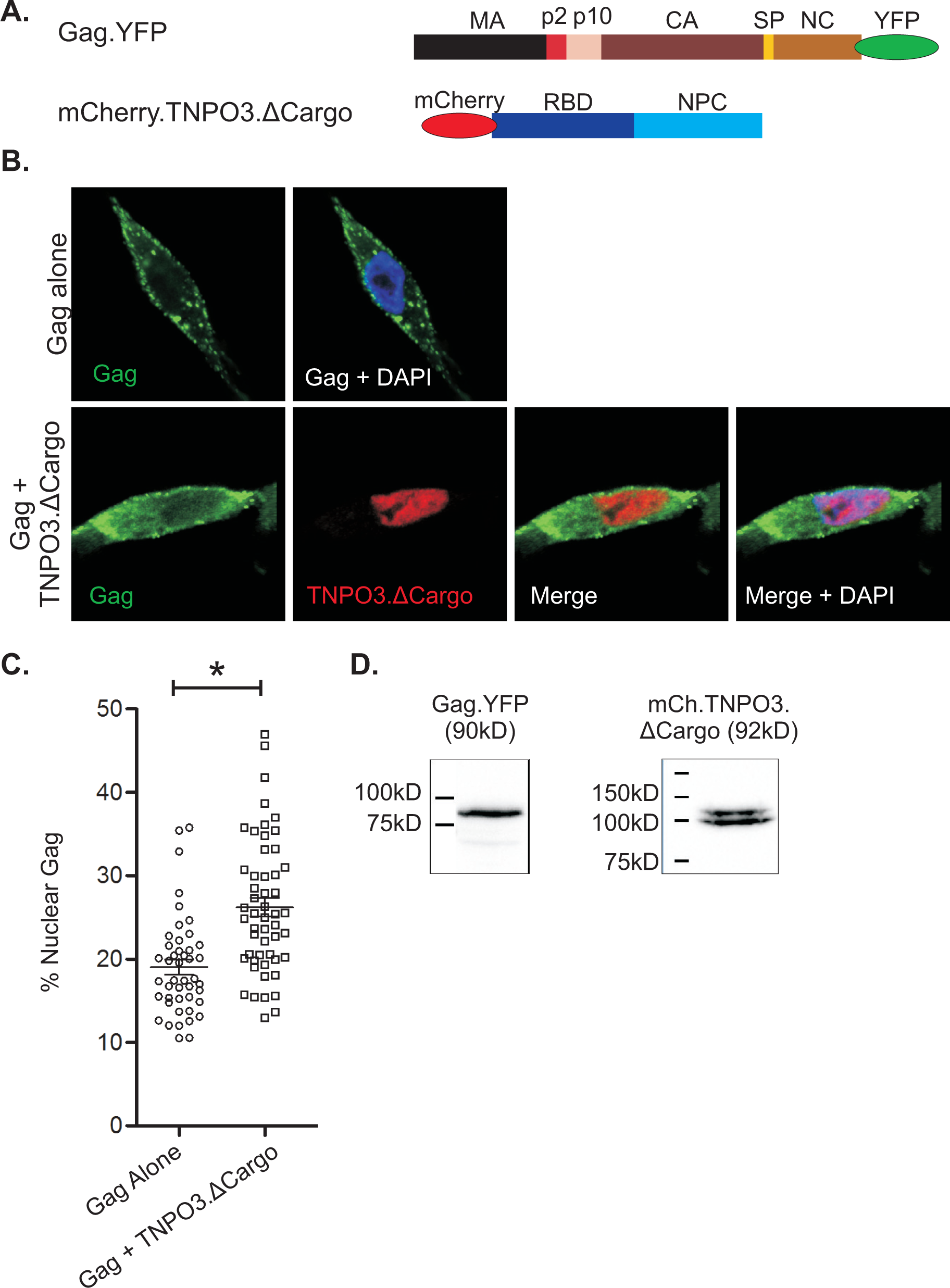
Gag nuclear accumulation with TNPO3 mutants. (A) Schematic representation of the constructs used, including Gag containing a C-terminal YFP fusion and TNPO3.ΔCargo, in which the entire cargo domain and a small portion of the NPC domain was deleted. (B) Top row shows Gag.YFP expressed alone in QT6 cells. The bottom row displays the accumulation of Gag.YFP with co-expression of mCherry.TNPO3.ΔCargo. The image chosen is representative of the mean nuclear flouorescence intensity of the population examined. (C) A scatter diagram plotting the percentage of nuclear Gag in each expressing Gag alone or co-expressed with TNPO3.ΔCargo indicates a significant increase in the nuclear population of Gag. At least 43 cells were analyzed from at least 3 independent experiments for each condition. The mean and standard error of the mean are shown with analysis using an unpaired Student’s t-test. The asterisk signifies p < 0.0001. Only cells expressing both Gag and mCherry.TNPO3.ΔCargo were analyzed. (D) Western blot detecting RSV Gag.YFP using an α-RSV antibody and mCherry.TNPO3.ΔCargo, which appears as a doublet, using an α-mCherry antibody indicate that proteins of the expected size were expressed.

To determine whether there was direct binding of Gag to TNPO3, in vitro affinity-tagged purification was performed with recombinant Gag, GST-tagged TNPO3, and deletion mutants of each, which were were expressed in *E. coli* and purified (Fig. 3A). RSV Gag was incubated with GST-TNPO3, and protein complexes were affinity purified using GST beads, separated by SDS-PAGE, and detected by western blot. Wild-type Gag was strongly associated with GST-TNPO3 but not with the GST protein alone (Fig. 3B). To determine whether the Gag-TNPO3 interaction was dependent on the presence of the MA domain, a Gag mutant containing a deletion of the first 82 amino acids of MA (ΔMBD.Gag) was tested (5, 42). ΔMBD.Gag showed reduced binding to GST-TNPO3 compared to full-length Gag (Fig. 3B). Additional Gag mutants were used to further define Gag-TNPO3 interaction sites, including a truncation at the end of the CA domain (Gag.ΔSPΔNC), a deletion of the C-terminal domain of CA through the end of NC (MA.p2.p10.CA-NTD), and a mutant expressing only CA and NC (CA.NC). Each Gag mutant associated to a small extent with GST-TNPO3, albeit much less efficiently compared to full-length Gag (Fig. 3B).

**Figure 3:**
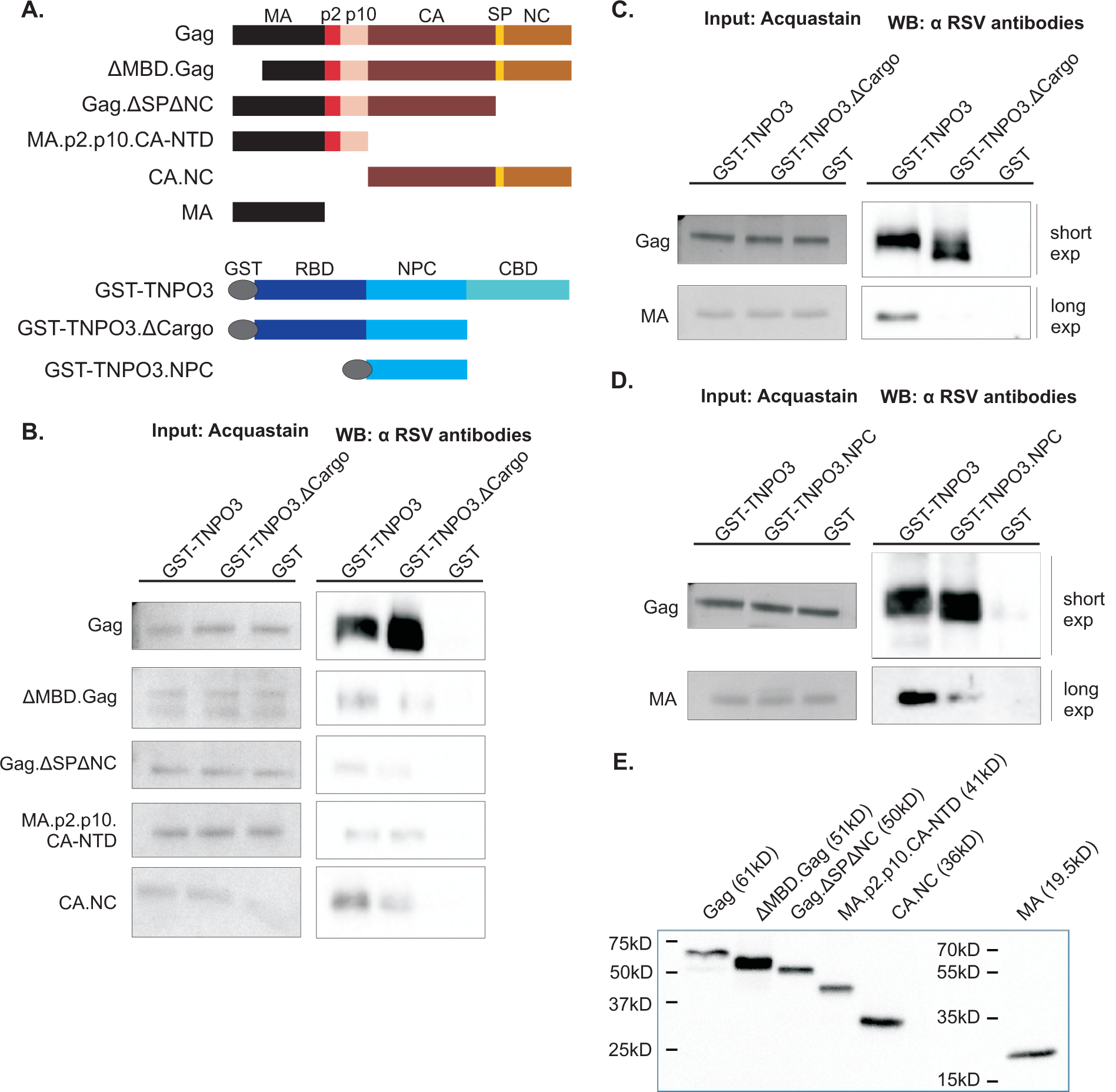
*In vitro* affinity-tagged purifications of Gag and TNPO3 protein complexes. (A) Schematic representation of constructs used (see Materials and Methods for details). CA-NTD contains the N-terminal domain of CA. GST was fused to the N terminus of TNPO3, with the ΔCargo mutant having the entire CBD and a small portion of the NPC domain deleted. The GST-TNPO3.NPC construct contains most of the NPC domain and a portion of the RanGTP binding domain. (B) In the affinity purifications, various recombinant Gag proteins were incubated with GST-TNPO3, GST-TNPO3.ΔCargo, or GST alone. In the left panel, one-tenth of the volume of the assay mixture was removed before the incubation step, separated by SDS-PAGE gel, and stained with Acquastain to directly visualize the Gag proteins put into into the binding assay (Input). In the right panel, the bound protein elutes were separated by SDS-PAGE and analyzed by western blot using α-RSV antibodies. (C and D) The same assay was used as in (B). Gag was detected in the linear range using a short exposure, whereas detection of MA required a longer exposure. At least 3 independent GST-tagged purifications were performed for each condition and a representative image is shown. (E) Western blot detecting purified Gag proteins using an α-RSV antibody to show full-length proteins of the expected molecular weights.

Next, we examined which domain of TNPO3 was required for binding to Gag. We utilized the mutant that lacked the CBD (GST-TNPO3.ΔCargo; Fig. 3A). We incubated TNPO3.ΔCargo with wild-type Gag, and found that Gag bound very strongly to TNPO3.ΔCargo (Fig. 3B, C). By contrast, there were only very weak interactions between TNPO3.ΔCargo and each of the Gag truncation mutants (Fig. 3B). A mutant of TNPO3 containing the NPC binding domain in the central region of TNPO3 (GST-TNPO3.NPC; Fig. 3A) was also tested for binding to Gag. As with full-length TNPO3 and the TNPO3.ΔCargo mutant, Gag bound strongly to the TNPO3.NPC protein (Fig. 3C).

To determine whether the mature MA protein was sufficient to mediate binding of full-length TNPO3 or the TNPO3 deletion mutants, similar GST-affinity pulldowns were performed. The MA protein bound full-length TNPO3, although much less efficiently compared to full-length Gag, with little to no binding to TNPO3.ΔCargo (Fig. 3C) or TNPO3.NPC (Fig. 3D). These results suggest that the requirement for robust binding requires the full-length Gag protein interacting with a TNPO3 construct that contains the NPC binding domain, but the CBD is dispensable. In addition, we conclude that the mature MA protein nor any of the Gag deletion mutants efficiently bind TNPO3, suggesting that the conformation of the intact Gag protein is required to mediate binding to TNPO3. Figure 3E demonstrates that that the purified Gag proteins used were single populations of the expected sizes as analyzed by western blot analysis.

To explore the interrelationship between TNPO3 and the other host factors involved in Gag nuclear import, the TNPO3 overexpression experiments were expanded to include other karyopherins known to interact with the Gag MA and NC NLSs. We predicted that overexpression of two import factors that bind distinct Gag NLSs would cooperate, resulting in enhanced import of Gag into the nucleus. Conversely, overexpression of two import factors that bind the same NLS could compete for binding sites on Gag, resulting in no further increase in nuclear entry. To test this hypothesis, each individual import factor was co-expressed with Gag, and the percentage of Gag in the nucleus increased significantly, as shown previously: co-expression with TNPO3 increased nuclear Gag from 15% to 21%; Imp11 co-expression increased nuclear Gag from 15% to 21%; and the percentage of nuclear Gag increased from 15% to 25% with Impβ co-expression (each p < 0.0001; Fig. 4A, C). When TNPO3 and Imp11 were co-expressed with Gag, the amount of nuclear Gag increased to 21%, which was not significantly different compared to TNPO3 or Imp11 expression alone. However, when TNPO3 and Impβ were both co-expressed with Gag, there was an increase in nuclear Gag from 21% (TNPO3 alone) to 28% (TNPO3 + Impβ; p < 0.0001) (Fig. 4B, C). These data demonstrate that Impβ and TNPO3 enhanced nuclear entry of Gag in an additive fashion, suggesting they have different binding sites in the MA region of Gag. By contrast, TNPO3 and Imp11 may compete for the same NLS in the MA domain of Gag and therefore are unable to increase the amount of Gag nuclear import when co-expressed.

**Figure 4:**
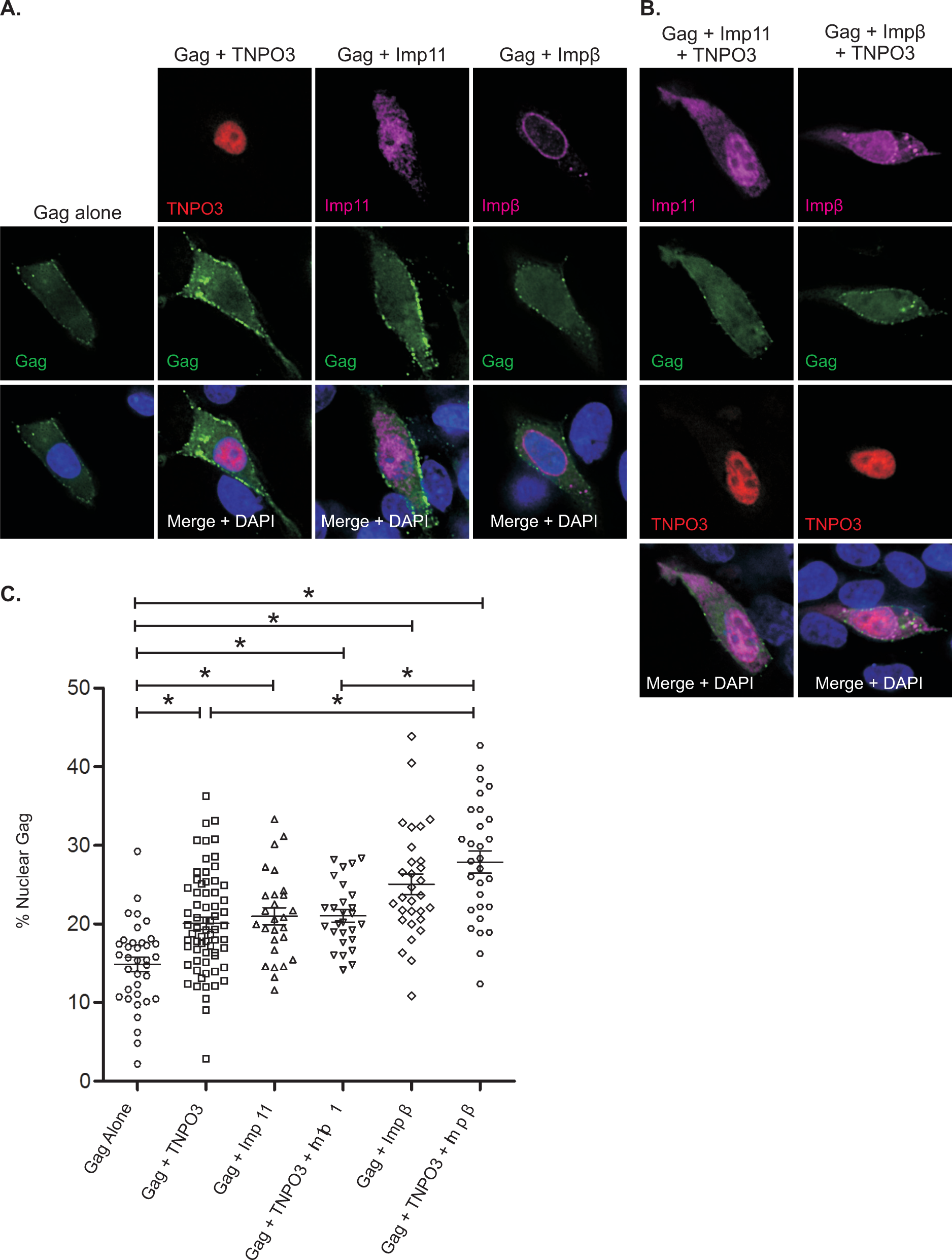
Effects of multiple import factors on Gag nuclear accumulation. (A) Visualization of subcellular localization of Gag.YFP (green) alone (first column) and with co-expression of individual import factors, mCherry.TNPO3 (red: second column), HA-importin-11 (magenta: third column), and HA-importin-β (magenta: fourth column). (B) Visualization of subcellular distribution of Gag.YFP with co-expression of importin-11 and TNPO3 (left column), and importin-β and TNPO3 (right column). (C) Scatter plot displaying the percentage of nuclear Gag alone, co-expressed with TNPO3, importin-11 or importin-β individually, and then co-expressed with the combination of TNPO3 + importin-11 or TNPO3 + importin-β. At least 27 cells were analyzed from two independent experiments for each condition. Group comparisons were analyzed using unpaired Student’s t-tests. The mean and the standard error of the mean are represented and an asterisk signifies statistically significant difference (p < 0.0001) between each compared group. For conditions of Gag co-expressed with an import factor, only cells expressing both tagged proteins (Gag and the import factors) were analyzed, and the images shown are representative of the mean fluorescence intensity of the population.

## Discussion

Nuclear trafficking of RSV Gag is important for efficient encapsidation of the viral genome (13, 42). To accomplish this vital step in virus replication, Gag uses multiple NLSs and host import pathways to gain entry into the nucleus. Previous studies revealed that Impα/β binds directly to a classical NLS in the NC domain of RSV Gag whereas Imp11 interacts directly with a noncanonical NLS in the MA domain of RSV Gag (14). Although we had previously identified TNPO3 (Mtr10p) as mediating MA nuclear entry in yeast (5), we wondered whether this finding would be verified in avian cells higher eukaryotic cells given that the MA sequence does not contain an RS-rich motif or a polyalanine domain characteristic of known TNPO3 cargoes (21, 22). In this report, we found that there is indeed a direct binding event between TNPO3 and RSV Gag that is is functionally relevant for the nuclear entry of RSV Gag in cellular import studies.

We demonstrated that overexpression of TNPO3 resulted in increased accumulation of Gag in the nucleus (Fig. 1) in an MA-dependent fashion. In cells, the normal function of TNPO3 is to import splicing factors into the nucleus through direct interactions between the C-terminal cargo-binding domain (CBD) and the RS or polyalanine domain of splicing factor cargoes (22, 23). We anticipated that Gag would bind the cargo domain of TNPO3, being a potential cargo itself; however, we found that the CBD was dispensable (TNPO3.ΔCargo) for the nuclear import of Gag based on the finding that TNPO3.ΔCargo expression in cells led to a significant increase in nuclear localization of Gag.

Our studies demonstrated that Gag binds directly to full length TNPO3 and the isolated NPC domain (Figure 3D), yet the interaction in vitro did not require the CBD (Figure 3C), which was consistent with the nuclear import data obtained in cells (Fig. 2). Finding that the TNPO3 CBD was dispensable for binding to Gag was somewhat surprising because nuclear import of splicing factors including SR proteins ASF/SF2, SC35, and CPSF6 are dependent on the TNPO3 CBD for import (17, 23). However, a recent study demonstrated that the N-terminus of TNPO3 (containing the RanGTP binding domain and part of the NPC interaction domain) interacts with the catalytic core domain and C-terminal domain of HIV-1 integrase, suggesting that other retroviral cargoes of TNPO3 bind to sites outside of the CBD (51). Based on our binding data, we conclude that RSV Gag binds primarily to the NPC domain of TNPO3 rather than to the CBD. The structural basis of this interaction will be interesting to explore in future studies.

We also found that the mature MA domain of Gag only bound to TNPO3 weakly in vitro, yet full length Gag bound TNPO3 strongly. Taken together with the in vivo experiments showing that MA is required for TNPO3-mediated import of Gag, we propose that MA must be present in the context of the full length Gag polyprotein to bind TNPO3. Interestingly, there was a small amount of TNPO3 pulled down with all of the deletion mutants of Gag tested in Fig. 3B, suggesting that regions of Gag in addition to MA contribute to the interaction with TNPO3. This finding is consistent with the structure of TNPO3, which is made of multiple HEAT repeats that encircle cargo (31), implying that multiple points of contact are important for TNPO3-Gag binding.

In previous studies, we examined RSV Gag nuclear import mediated by one import factor at a time, even though Gag contains two separate NLSs and binds three diffierent karyopherins (5, 14). Therefore, we wondered whether there was cooperation or competition between different host import factors responsible for Gag nuclear entry. Data presented in Fig. 4 indicates that TNPO3 can cooperate with Impβ but not Imp11 to drive Gag into the nucleus. This finding implies that factors interacting with different NLSs of Gag (i.e., TNPO3 with MA and Impα/Impβ with NC) are able to function in an additive manner to mediate Gag nuclear import. These findings also suggest that factors binding to the same domain of Gag (i.e. TNPO3 and Imp11 both bind to MA) cannot cooperate for Gag nuclear import, potentially because they bind to the same site or to sites in close proximity to one another.

A remaining question is why RSV Gag utilizes three different nuclear import pathways. One possibility is that each import pathway is used for Gag to reach a particular location in the nucleus or perform a specialized function. For example, Impβ transports a variety of proteins into the nucleus including transcription factors, cell cycle regulators, histones, and ribosomal proteins [reviewed in (6)]. Imp11 transports ribosomal proteins and E2-ubiquitin-conjugating enzymes [reviewed in (6)]. TNPO3 transports SR splicing factors to splicing speckles (22-24), which are adjacent to transcription sites (48). Splicing factors are activated through phosphorylation, triggering them to traffic to sites of active transcription to process newly transcribing RNAs (33, 37), with rapid exchange of splicing factors between sites of transcription and speckles (20, 50). We previously showed that a nuclear-localized mutant of RSV Gag (Gag.L219A) has a high degree of colocalization with the phosphorylated forms of SR protein splicing factors SF2 and SC35 (40). Thus, it is possible that Gag enters the nucleus using the TNPO3 pathway to reach nuclear speckles adjacent to sites of nascent mRNA transcription (17, 50). Further experiments will need to be performed to dissect the precise role of TNPO3 in Gag nuclear trafficking and RSV biology.

## Materials and Methods

### Cells and Plasmids

QT6 quail fibroblast cells were maintained as described in (8) and were transfected with the calcium phosphate precipitation method.

#### Tissue culture expression

Gag.GFP was described in (42). Gag.ΔMA_5-86_.GFP and Gag.ΔMA_5-148_.GFP were described previously (5). Gag.YFP and Gag.ΔNC.YFP were previously described (18). GFP-TNPO3 and GFP-TNPO3.ΔCargo (29) were gifts of Nathaniel Landau (NYU Langone Medical Center). Ceruleun.TNPO3.CBD was a generous gift from Yaron Shav-Tal (Bar Ilan University). mCherry.TNPO3 was created by PCR amplifying the TNPO3 coding sequence from GFP.TNPO3 with flanking *Xho*I and *Sal*I restriction sites and inserting into mCherry.N2 digested with those same enzymes. pKH3.Importin 11 and pKH3.Importin β, encoding HA.Importin 11 and HA.Importin β, respectively, were described in (14).

#### E. coli protein expression

pET28.TEV-Gag.3h encoding RSV Gag with an N-terminal cleavable 6-histidine tag (Gag) is described in (14). Gag.ΔNC.ΔSP was created by inserting two consecutive, in-frame stop codons into pET28.TEV.Gag.3h preceding the SP coding sequence. Gag.ΔMBD was created by PCR amplification of the Gag coding region starting at amino acid 83 and terminating at the N-terminus of NC with the insertion of stop codons. This product was ligated into appropriately digested pET24a+. CA.NC was created by PCR amplification of CA and NC inserting a NdeI site at the N-terminus and a HindIII site at the C-terminus. The amplicon was inserted into the digested pET28(-His).Gag.ΔPR using the same sites. MA.p2.p10.CA-NTD was created similarly as CA.NC with the NdeI site at the N-terminus of MA and two STOP codons followed by the HindIII site after the 444 nt of CA. MA was described previously (14). pGEX6P3.hTNPO3 (21) encoding GST.TNPO3 for bacterial expression was a gift from Alan Engelman (Dana Farber Cancer Institute). GST.TNPO3.ΔCargo_1-501_ was created from pGEX6P3.hTNPO3 by inserting three in frame, consecutive stop codons following the sequence encoding amino acid 501 of TNPO3 by Quikchange PCR mutagenesis (28). GST.TNPO3.NPC was created by deleting most of the Ran binding domain using primers: 5’ – GGA GAA AAC CTT TAC TTC CAG GG-3’ and 5’ – AAT GGA TCC CAG GGG CCC-3’ using the Q5 Site-Directed Mutagenesis protocol according to the manufacturer guidelines (New England Biolabs). Michael Malim (King’s College London) kindly provided the bacterial expression construct encoding GST-Importin β.

### Cell fixation and immunofluorescence

QT6 cells were grown on glass coverslips and fixed with 2% paraformaldehyde (PFA) in phosphate buffered saline (PBS) (supplemented with 5 mM EGTA and 4 mM MgCl_2_, and adjusted to pH 7.2-7.4 with HCl) or 3.7% PFA in 2x PHEM buffer (3.6% PIPES, 1.3% HEPES, 0.76%EGTA, 0.198% MgSO4, pH to 7.0 with 10M KOH) (32) for 15 minutes at room temperature. Cells expressing HA.Importin β or 11 were then permeabilized with ice cold 100% methanol for 2 minutes on ice and blocked in 5% goat serum (Rockland) diluted into PBS for at least 2 hours at room temperature. Cells were stained with mouse anti-HA antibody (Genscript) diluted 1:500 in PBS supplemented with 0.5% goat serum and 0.01% Tween-20 (Sigma) for at least one hour in a humidified chamber.

After primary antibody incubation, the cells were incubated with goat anti-mouse antibody conjugated to Cy5 (Molecular Probes) diluted 1:500 or Alexa 647 (Life Technologies) diluted 1:1000 in PBS for 1 hour at room temperature. Cells were stained with DAPI at 5 μg/ml and mounted with Slow-Fade mounting medium (Molecular Probes) or Prolong Diamond (Thermo Fisher Scientific).

### Microscopy

Images were captured either using a DeltaVision DV Elite (Applied Precision) wide field deconvolution microscope with a 60x oil immersion objective or a Leica SP8 TCS scanning confocal microscope equipped with a 63X oil immersion objective and White Light Laser (WLL). For the DeltaVision, 0.2 μm optical slices encompassing the entire cell were captured and deconvolved with softWoRx 5.0 (Applied Precision). From the deconvolved image stack, a single slice encompassing the widest section of the nucleus was exported for each channel as an uncompressed TIFF file for subsequent display and analysis. For images acquired using the SP8, we employed sequential scanning between frames, averaging four frames per image. The 405 nm UV laser was used to image the DAPI signal 10% laser power with an emission detection window of 415 – 466 nm using the PMT detector. GFP was imaged using the WLL excited with the 489 nm laser line and a hybrid detector window of 495 – 559 nm. YFP was imaged using the WLL with a laser line excitation of 514 nm and a hybrid detector window of 519 – 569 nm. mCherry was imaged using the WLL with a laser line excitation of 587 nm and a hybrid detector window of 592 – 656 nm. All channels using the hybrid detectors had a time gating of 0.3 to 6.0 ns.

### Quantitation of nuclear localization

ImageJ software version 1.46m was used to analyze cells for the amount of Gag present in the nucleus, as determined by fluorescence signal (1). The sum of all the pixel intensities in the Gag channel, expressed in ImageJ as the “Integrated Density,” for a region encompassing the entire cell was divided by the integrated density of the nucleus to calculate the percentage of the total cellular Gag pool residing in the nucleus. Outliers with a *p* value less than 0.05 as determined by Grubbs’ test (GraphPad Software Inc, <http://graphpad.com/quickcalcs/Grubbs1.cfm>) were removed from subsequent analyses. GraphPad Prism 5 (GraphPad Software, Inc.) was used to create all graphs, perform linear regression analysis, calculate the mean, and determine *p* values. *p* values were calculated by unpaired *t*-test as only pairs (i.e. no more than two) of conditions are analyzed simultaneously.

### Expression and purification of recombinant RSV Gag proteins

All constructs for protein expression and purification were transformed into BL21 Gold DE3 pRIL *E. coli*. The purity of all protein preps was verified with Coomassie staining following SDS-PAGE and/or western blot analysis with appropriate antibodies. All sonications were performed on ice with an S-4000 sonicator (Misonix, Inc.) using a ½” tip. All Gag constructs were expressed in ZYP-5052 autoinduction medium, lysed in BugBuster Primary Amine Free Protein Extraction Reagent (Novagen) supplemented with recombinant lysozyme. PEI was added to a final concentration of 0.15% and lysate was centrifuged at 21,000 RCF for 30 minute to remove cell debris. The protein remaining in the soluble fraction was precipitated for 30 minute at room temperature with concentrated ammonium sulfate. The pellet containing the Gag protein was resuspended and clarified by centrifugation prior to chromatographic separation and elution with a sulfopropyl cation exchange column. Peak eluted fractions were dialyzed against 25 mM HEPES pH 7.5, 500 mM NaCl, 0.1 mM EDTA, 0.1 mM TCEP, and 0.01 mM ZnSO_4_ prior to concentration, aliquoting, and storage at -80°C.

### Expression and purification of recombinant GST-tagged proteins

Purified GST protein was a gift from John Flanagan (Penn State College of Medicine). GST.TNPO3 was grown in two 250 ml cultures of ZYP-5052 supplemented with ampicillin and incubated at 37°C for 19.5 hours. Cell pellets were harvested by centrifugation and stored at -20°C prior to purification. A batch purification protocol adapted from (15) was used to purify GST.TNPO3. Cell pellets were thawed on ice, homogenized into PBS containing Roche Complete EDTA free protease inhibitors (Roche). Ready-lyse lysozyme and Omnicleave nuclease (Epicentre) were added and the mixture was permitted to rock on ice for 15 minutes. The homogenate was then sonicated three times at 80% power, with a one-minute recovery between each sonication. Lysate was clarified for 30 min at 21,000 RCF at 4°C and passed through a 0.45 μm filter. The soluble portion was incubated with gentle end over end mixing for 3 hours at 4°C with Glutathione Sepharose 4 Fast Flow Beads (GE) prewashed with PBS. Following binding, the beads were washed three times with PBS to remove unbound proteins. Bound proteins were eluted with a 30 minute incubation at 4°C with Elution Buffer (50 mM Tris pH 8.0, 40 mM reduced glutathione, and Roche Complete protease inhibitors). At the end of the incubation, beads were pelleted and the supernatant removed to a prechilled tube. Two additional elution steps were performed. All elutions were pooled and dialyzed against TNPO3 storage buffer (50 mM HEPES pH 7.4, 150 mM NaCl, 10% Glycerol, and 2 mM DTT). Following dialysis, purified GST.TNPO3 was concentrated with an Amicon Ultra Centrifugal Filter Device (Millipore), aliquoted, and stored at -80°C prior to use. GST.TNPO3.ΔCargo and GST.TNPO3.NPC were expressed at 30°C, but otherwise expressed and purified identically to GST.TNPO3.

### In vitro GST affinity purification protein-protein interactions assays

The protocol for GST affinity purification assays was adapted from (21). All proteins used in purification assays were performed at equimolar of 185 nM in 540 μl pull-down buffer (150 mM NaCl, 5 mM MgCl_2_, 5 mM DTT, 0.1% NP-40, 25 mM Tris-Cl pH 7.4). For the input gel, 40 μl was removed. Proteins were incubated for 1 hour at room temperature with gentle end over end rotation. 60 μl of a 50% slurry of glutathione beads (Glutathione Sepharose 4 Fast Flow, GE) prewashed four times in pull-down buffer were then added to the complexes and incubated for 2 hours at room temperature with gentle end over end rotation. The beads were pelleted by centrifugation at 800x*g* for 2 minutes. The supernatant containing unbound proteins was removed and the beads containing the bound protein complexes were washed four times with 10 packed bead volumes of pull down buffer. Following the final wash, one packed bead volume of elution buffer (25 mM Tris-Cl and 40 mM reduced glutathione, pH 8) was added, mixed with the beads, and placed on ice for 10 minutes to elute the bound complexes. Following elution, beads were pelleted by centrifugation at 800x*g* for 2 minutes at 4°C. The supernatant containing bound complexes was then removed to a clean microcentrifuge tube and 4x SDS-PAGE loading buffer (250 mM Tris-HCl, pH 6.8, 40% glycerol, 0.4% bromophenol blue, 8% SDS, and 8% β-mercaptoethanol) was added to 1x final concentration. The samples were then heated for 5 minutes at 85°C and separated by SDS-PAGE and analyzed by Western blot using rabbit anti-GST antibody (Genscript), rabbit α-RSV CA, rabbit α-RSV MA.p2, rabbit α-RSV MA.MDB, rabbit α-RSV polyclonal antibody and HRP-conjugated secondary antibodies (Invitrogen). RSV antibodies were generous gifts from Rebecca Craven (Penn State College of Medicine). The input gels were visualized with Acquastain dye (Bulldog Bio). mCherry tagged proteins were analyzed by Western blot using mouse anti-mCherry (Abcam ab125096).

## Acknowledgments

We would like to thank the following scientists for their generosity in supplying reagents, Dr. Nathaniel Landan (NYU Langone Medical Center), Dr. Alan Engelman (Dana Farber Cancer Institute), Dr. Michael Malim (King’s College London), Dr. Yaron Shav-Tal (Bar Ilan University), Dr. John Flanagan and Dr. Rebecca Craven (PSU College of Medicine). Special thanks to Malgorzata Sudol in the Department of Medicine at the Penn State College of Medicine for technical assistance. We would like to acknowledge the Microscopy Imaging Core Facility at PSU College of Medicine for use of the confocal [Leica SP8-1S10OD010756-01A1 (CB)] and deconvolution microscopes and the Imaris (Bitplane imaging analysis software. This project was funded in part by NIH R01 CA076534 (LJP) and F31 CA196292 (BLR).

## Notes

### Competing Interest Statement

The authors have declared no competing interest.

